# Sex-Specific Induction of H3K27me1 in the Prefrontal Cortex Mediates the Enduring Effects of Early Life Stress

**DOI:** 10.1101/2025.11.25.690167

**Authors:** Jonathan Nguyen, Nidhi Byragoni, Nneka Arinzeh, Alessandro Bortolami, Simone Sidoli, Scott J. Russo, Eric J. Nestler, Angélica Torres-Berrío

## Abstract

Early life stress (ELS) is a strong risk factor for several neurodevelopmental and psychiatric disorders and is associated with persistent molecular alterations in corticolimbic regions, including the prefrontal cortex (PFC). Here, we combined unbiased proteomics profiling and viral-mediated gene transfer to characterize the enduring epigenetic changes in the PFC of male and female mice previously exposed to ELS. We found that ELS induced sex-specific and age-dependent accumulation of monomethylation of lysine 27 at histone H3 (H3K27me1), with a transient increase in adolescent females, and a delayed but enduring effect in adult males. ELS-induced H3K27me1 accumulation in the PFC involved changes of SUZ12, a subunit of the polycomb repressive complex 2 (PRC2), which controls H3K27 methylation patterns. Indeed, expression of the catalytic domain of SUZ12, VEFS, in PFC neurons during adolescence led to sex differences in social and cognitive alterations in adulthood. Together, our results demonstrate that H3K27me1 functions as a “chromatin scar” in the PFC, where it mediates lifelong susceptibility to ELS in a sex-specific manner.

## INTRODUCTION

Early life is a critical period of brain plasticity and is highly sensitive to adverse experiences such as stress exposure, which can shape mental health trajectories across the lifespan^1–3^. Early life stress (ELS), whether in the form of child maltreatment, parent neglect, undernutrition, or sexual abuse, affects millions of children and adolescents around the world each year, increasing the risk for depression and other stress-related disorders by two- to fourfold^2, 4–8^. Most recently, disruptions in daily routines, social isolation, and quarantine during the COVID-19 pandemic were severe stressors worldwide^9, 10^, which triggered large increases in depression and anxiety symptomatology^11, 12^, including suicidal behaviors among middle- and high-school students^12–14^. These data indicate that a mental health crisis within the youth population is unfolding in real-time and highlights the urgent need for understanding the molecular mechanisms underlying the enduring effects of stress exposure during sensitive periods of neurodevelopment^1, 3, 15, 16^.

The effects of ELS are associated with persistent transcriptional, cellular, and circuit adaptations across the corticolimbic pathway, including the prefrontal cortex (PFC) and nucleus accumbens (NAc)^1, 3, 17–20^, two key regions that play a critical role in mood regulation and cognitive control^1, 21^. Indeed, genome-wide RNA sequencing (RNA-seq) studies demonstrate that ELS leads to broad transcriptional disruptions in the PFC and NAc that sensitize stress responses later in life^17, 22, 23^. These effects are linked to epigenetic alterations, including changes in histone posttranslational modifications^18, 19, 24^ or chromatin accessibility^22^, which work cooperatively to promote or repress future transcriptional states^17, 25^.

Recent work from our group identified the histone mark, H3K27me1 (monomethylation of Lys 27 of histone H3), as a “chromatin scar*”* that confers lifelong susceptibility to stress^19^. Specifically, we found that mice exposed to early life or adult stress displayed increased abundance of H3K27me1 in the NAc. This effect was observed only in adult mice that are susceptible to stress, with resilient mice not displaying H3K27me1 induction in the NAc. Increased levels of H3K27me1 in the NAc were specific for medium spiny neurons (MSNs) expressing the dopamine receptor 1 (D1), a critical cell type within the NAc that mediates the development of susceptible vs resilient stress-related phenotypes^26, 27^. Our results also showed that H3K27me1 accumulation in the NAc was male-specific and involved SUZ12, a core protein of the polycomb repressive complex-2^19^ (PRC2), which controls H3K27 methylation patterns^28–33^. A question that remains is whether ELS also induces sex-specific epigenetic programs in the PFC and, in turn, prompts divergent behavioral trajectories between male and female mice. We focused on the PFC because of its strong influences on NAc function and because this region undergoes a protracted maturational period during adolescence that continues through adulthood^1, 34–36^.

Here, we exposed male and female pups to ELS, by combining maternal separation and reduced nestling and bedding^17–19, 25^, to characterize sex-specific histone profiles in the PFC during adolescence and adulthood. We found that ELS induces H3K27me1 accumulation in the PFC but, unlike in the NAc^19^, this effect occurs in an age-dependent and sex-specific manner, with a transient increase in the PFC of adolescent females, and a delayed but enduring increase in the PFC of adult males. ELS-induced H3K27me1 accumulation in the PFC, like in the NAc, involves changes in SUZ12. Indeed, expression during adolescence of the VEFS-domain of SUZ12, the C-terminal domain that contributes to the catalytic actions of PRC2, leads to sex-specific social and cognitive alterations in adulthood, recapitulating some of the behavioral alterations induced by ELS. This study thereby establishes a novel function of H3K27me1 across the corticolimbic pathway and demonstrates its role as an important chromatin scar that mediates the lifelong effects of ELS.

## RESULTS

### ELS leads to sex-specific behavioral alterations in adult mice

Our previous work established that ELS, in the form of maternal separation and reduced bedding during postnatal day (PND) 10 to PND 17, increases the susceptibility of male and female mice to develop certain depression- and anxiety-like behavioral abnormalities in response to stress during adulthood^17–19, 25^. To establish whether ELS also induces changes in cognitive function in adulthood and whether these effects are also exacerbated by a second stressor in adulthood, we subjected male and female pups to our ELS paradigm (**Figure 1A**). Standard-raised pups (Std) stayed with the dams and were handled during the same period. Once all mice reached adulthood, a subset was subjected to subthreshold social defeat stress (SbD), a paradigm that induces a rapid stress response but does not induce social avoidance to a CD1 aggressor mouse^37^. As expected, male and female mice subjected to ELS followed by adult exposure to SbD (ELS-SbD) displayed reduced social interaction compared to Std (Std-CON), ELS-CON, or SbD alone (SbD-CON) conditions (**Figure 1B-C**), which is consistent with our previous work using a similar “double-hit stress” approach^17–19, 25^.

**Figure 1.**
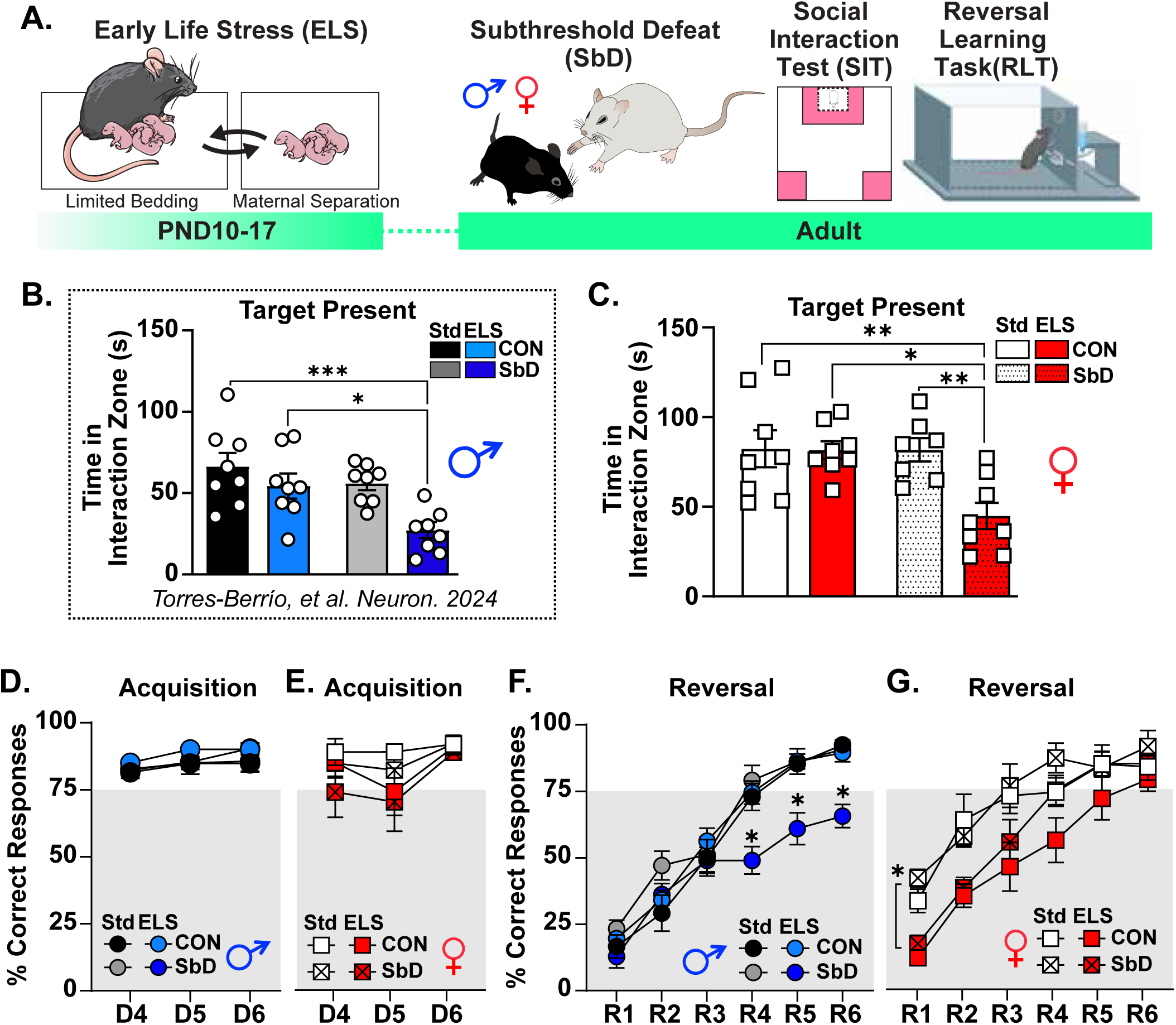
Effects of ELS and adult stress on behavioral outcomes. **(A)** Schematic of the “double-hit” stress procedure. Mice subjected to ELS further received a second stress exposure, subthreshold social defeat (SbD), in adulthood. **(B)** Social interaction in male mice exposed to this double-hit stress. Adapted from^19^. **(C)** Social interaction in female mice exposed to double-hit stress: Two-way ANOVA: ELS: F_(1,27)_=6.05, p=0.02; SbD: F_(1,27)_=5.62, p=0.025; ELS by SbD: F_(1,27)_= 5.96, p=0.021. Tukey’s comparison: ELS-SbD different from Std-CON (*p<0.05), ELS-CON (**p<0.01), and Std-SbD (**p<0.01). **(D)** Percentage of correct lever presses during the last three acquisition sessions in male mice: Three-way ANOVA: ELS effect: F_(1,21)_=0.80, p=0.38; SbD effect: F_(1,21)_=1.63, p=0.21; Session effect: F_(2,21)_=2.445, p=0.11; ELS by SbD by Session interaction: F_(2,21)_=0.25, p=0.77. **(E)** Percentage of correct lever presses during the last three acquisition sessions in female mice: Three-way ANOVA: ELS effect: F_(1,21)_=0.94, p=0.34; SbD effect: F_(1,21)_=7.53, p<0.05; Session effect: F_(2,21)_=2.445, p=0.11. ELS by SbD by Session interaction: F_(2,21)_= 0.091. **(F)** Combined exposure to ELS and SbD impaired reversal learning in male mice. Three-way ANOVA: ELS effect: F_(1,42)_=14.82, p<0.001; SbD effect: F_(1,42)_=6.61, p<0.05; session effect: F_(5,42)_=113, p<0.0001; ELS by SbD interaction: F_(1,42)_=31.39, p<0.001; ELS by session interaction: F_(5,42)_=4.03, p<0.05; SbD by session interaction: F_(5,42)_=2.09, p=0.08. Sidak’s comparison for ELS by SbD interaction: ELS-SbD different from Std-CON at R4, R5 and R6, *p<0.05. **(G)** ELS impaired reversal learning in female mice. Three-way ANOVA: ELS effect: F_(1,42)_=7.49, p<0.01; SbD effect: F_(1,42)_=38.01, p<0.0001; Session effect: F_(5,42)_=79.67, p<0.0001. Sidak’s comparison for ELS effect: ELS different from Std at R1, and R2, *p<0.05.

We next assessed experimental mice in the operant reversal learning task (RLT) to examine the effects of ELS on cognitive flexibility. The RLT task allows for the study of the ability to adjust behavior when the contingencies previously learned are reversed^38^, which is affected in mice susceptible to social stress^19^. For this, we trained Std-CON, ELS-CON, Std-CON, and ELS-SbD mice to press one of two levers for a 0.2% saccharin-water reward (acquisition phase)^19^. Daily trials were given until all mice met a learning criterion of 75% correct responses over three consecutive days, with no significant differences between groups identified during this phase (**Figure 1D-E**). In the reversal phase, all experimental mice were trained to press the previously unrewarded lever to obtain the same saccharin reward. Interestingly, we observed a sex-specific impairment of cognitive flexibility in mice exposed to ELS (**Figure 1F-G**). Indeed, Std-CON, ELS-CON, or Std-SbD male mice exhibited similar reversal, whereas those exposed to ELS followed by SbD had a lower percentage of correct responses during reversal days 4, 5, and 6 compared to Std-CON and ELS-CON, indicating impaired reversal learning in this group (**Figure 1F**). By contrast, female mice exposed to ELS, either alone or in combination with SbD, showed a lower percentage of correct responses during the first two days of reversal learning, but then achieved similar performance during the late stages of the task (**Figure 1G**). Overall, these findings indicate that, while male and female mice exposed to double-hit stress exhibit social interaction deficits, they display sex differences in cognitive flexibility, with females being more susceptible to the effects of ELS alone.

### ELS induces sex-specific methylation of H3K27 in the PFC

We recently showed that mice exposed to ELS^18, 19^ display dynamic alterations in histone modifications in the NAc^18, 19^. Therefore, we aimed here to characterize global changes in histone modifications induced by ELS in the PFC, a brain region that is also highly sensitive to the effects of stress^1, 21^ and directly involved in cognitive flexibility and top-down emotional regulation^35^. To this end, we used mass spectrometry, an unbiased proteomic approach that enables quantification of hundreds of histone marks^39^. PFC tissue, including prelimbic and infralimbic subregions, from male and female mice exposed to ELS and their Std-raised counterparts^18^ was collected during early adolescence at PND21 or in early adulthood at PND60^37^ and processed for mass spectrometry (**Figure 2A**). Analysis of the relative abundance of single peptides revealed a small subset of histone modifications induced by ELS exposure in the PFC (**Figure 2B, Figure S1A, Table S1**). Strikingly, we found age-dependent and sex-specific alterations of the methylation dynamics of K27 in the histone variant H3.3 (**Figure 2C**), the dominant form of H3 in adult brain neurons^24, 40, 41^. Indeed, there was a higher H3.3K27me1 abundance in adult male mice exposed to ELS (**Figure 2D**), and a corresponding decrease of H3.3K27me2 (**Figure 2E**), compared to Std-raised counterparts. By contrast, female mice exposed to ELS showed a transient increase of H3.3K27me1 in adolescence, but these changes normalized to Std-raised levels in adulthood (**Figure 2F**). ELS did not induce changes in levels of H3.3K27me2 in females (**Figure 2G**). These findings indicate that ELS alters the methylation dynamics of H3K27 in the PFC and that these effects occur in a sex-specific and age-dependent manner. Because H3.3K27me1 (hereafter referred to as H3K27me1) also accumulates in the NAc following ELS exposure^19^, these results suggest that changes in H3K27me1 across several corticolimbic brain regions may mediate stress susceptibility across the lifespan^19^.

**Figure 2.**
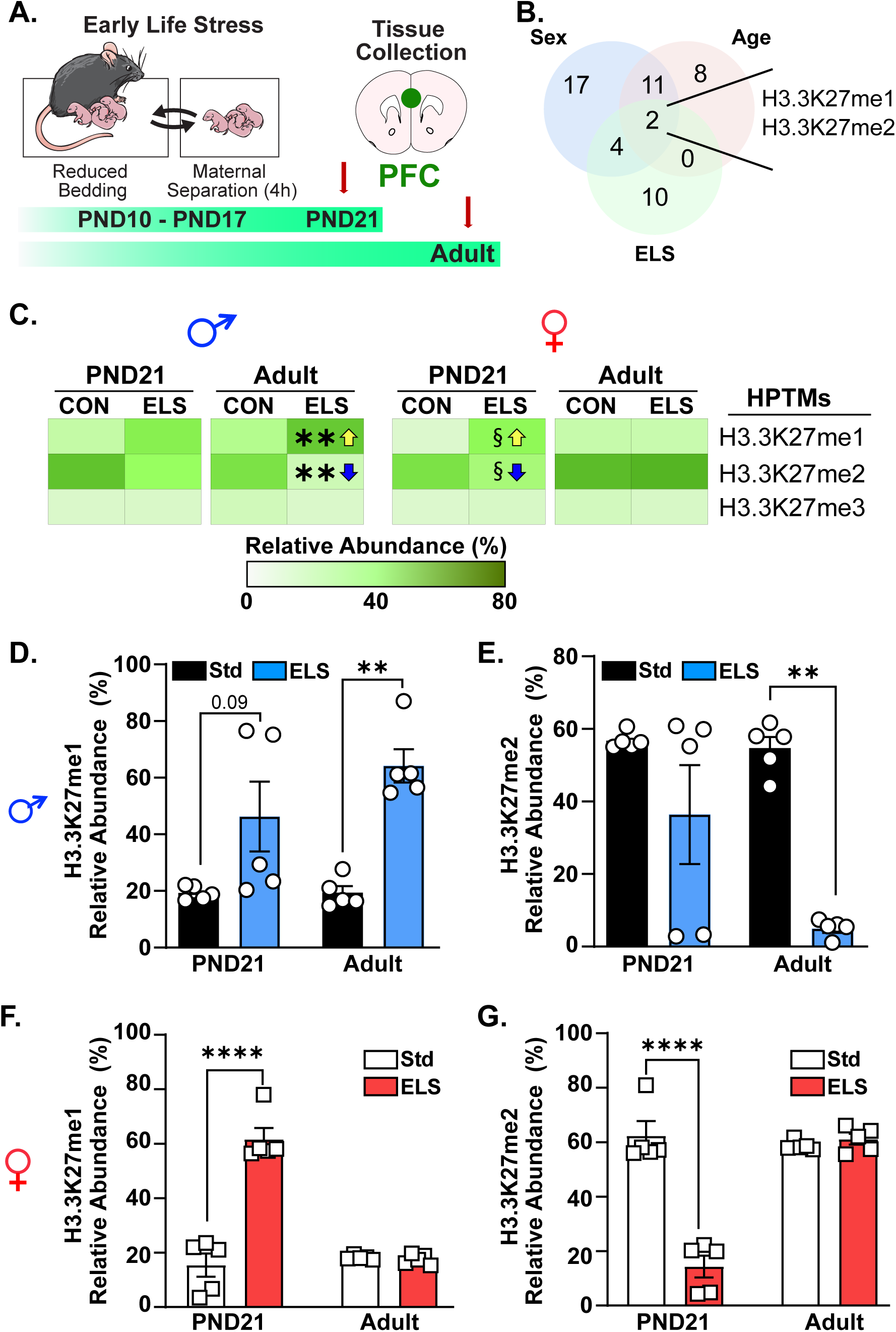
ELS induces sex-specific dynamic methylation of H3K27 in the PFC. **(A)** Schematic and timeline of ELS experiment and tissue collection (n= 5 samples from 3-4 different litters, 2 mice/group pooled tissue). **(B)** Venn diagrams showing significant H3.3K27me1 and H3.3K27me2 overlap according to Sex, Age and Group. Related to Supplementary Table 1. **(C)** Heatmap for relative abundance of H3.3K27 modifications for males and female mice. Arrows indicate significance (yellow, increase: **p<0.01; blue, decrease: **p<0.01). **(D)** H3.3K27me1 in males: Two-way ANOVA: ELS effect: F_(1,16)_=26.68, p<0.0001; Age effect: F_(1,16)_=1.66, p=0.22; ELS by Age interaction: F_(1,16)_=1.65, p=0.21. Sidak’s comparison for ELS effect: ELS different from Std in Adult, **p<0.01. **(E)** H3.3K27me2 in males: Two-way ANOVA: ELS effect: F_(1,14)_=21.89, p<0.001; Age effect: F_(1,14)_=4.02, p=0.06; ELS by Age interaction: F_(1,14)_=4.33, p=0.06. Sidak’s comparison for ELS effect: ELS different from Std in Adult, **p<0.01. **(F)** H3.3K27me1 in females: Two-way ANOVA: ELS effect: F_(1,16)_=58.01, p<0.0001; Age effect: F_(1,16)_=47.8, p<0.0001; ELS by Age interaction: F_(1,16)_=61.74, p<0.0001. Sidak’s comparison for ELS effect: ELS different from Std at PDN21, ****p<0.01. **(G)** H3.3K27me2 in females: Two-way ANOVA: ELS effect: F_(1,16)_=41.81, p<0.0001; Age effect: F_(1,16)_= 37.33, p<0.0001; ELS by Age interaction: F_(1,16)_=50.8, p<0.0001. Sidak’s comparison for ELS effect: ELS different from Std in Adult, ****p<0.01. Data: Mean ± SEM.

### ELS-induced increase in H3K27me1 occurs in PFC pyramidal neurons

Pyramidal excitatory neurons in the PFC are highly sensitive to the effects of stress^26, 27, 42, 43^. Therefore, we next assessed whether ELS-induced changes in H3K27me1 occur within these cell populations. For this, we exposed male mice to ELS or Std conditions and collected PFC tissue at PND60 (**Figure 3A**). Tissue was processed with antibodies that label H3K27me1, which recognize K27me1 in H3 and H3.3^19^, and SMI-32, which is a marker of cortical pyramidal neurons^43–46^. We observed that ELS-exposed mice exhibited increased H3K27me1 in PFC pyramidal neurons (**Figure 3C**). By contrast, no increase was apparent collectively across all other PFC cell types (not shown). Within the same mice we replicated our published finding^18^ that ELS increases H3K27me1 in D1-MSNs of the NAc (**Figure 3D-E**), with a trend seen for D2-MSNs (**Figure 3E**).

**Figure 3.**
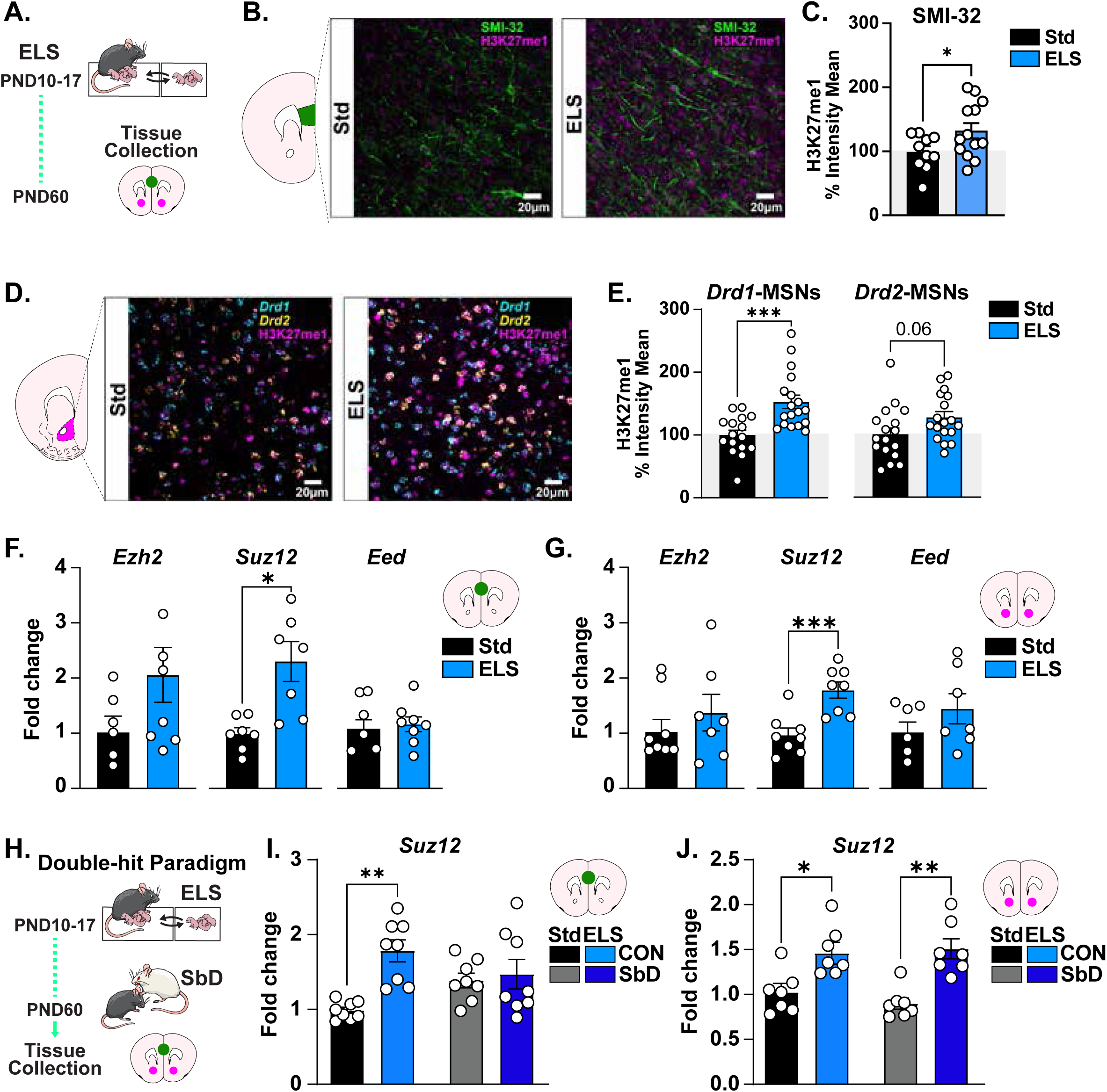
Cell-type and region-specific changes in H3K27me1 and PRC2 subunit expression. **(A)** Schematic and timeline of ELS experiment in male mice and tissue collection. **(B)** Coronal section showing H3K27me1 (magenta) in SMI-32 (green) pyramidal neurons (n= 2 bilateral sections; 3 mice/group). Scale bars: 20 μm. **(C)** Percentage of H3K27me1 fluorescence intensity in SMI-32 positive pyramidal neurons: t_(32)_=2.13, p<0.05. **(D)** Coronal section showing H3K27me1 (magenta) in *Drd1*- (cyan) and *Drd2*- (yellow) MSNs (n=3-4 bilateral sections; 4 mice/group). Scale bars: 20μm. **(E)** Percentage of H3K27me1 fluorescence intensity in *Drd1*-and *Drd2*-MSNs. *Drd1*-MSNs: t_(32)_=3.92, p<0.001. *Drd2*-MSNs: t_(33)_=2.01, p=0.06. **(F)** Gene expression of PRC2 subunits in the PFC (n=6-7/group). *Ezh2* mRNA: t_(13)_=1.25, p=0.23; *Suz12* mRNA: t_(13)_=2.36, p<0.05; *Eed* mRNA: t_(13)_=1.05, p=0.30. **(G)** Gene expression of PRC2 subunits in NAc (n=6-8/group). *Ezh2* mRNA: t_(13)_=0.67, p=0.51; *Suz12* mRNA: t_(14)_=4.19, p<0.001; *Eed* mRNA: t_(13)_=0.38, p=0.70. **(H)** Timeline of “double-hit” stress experiment (n=6-8/group). **(I)** *Suz12* mRNA in PFC: Two-way ANOVA: ELS effect: F_(1,28)_=10.87, p<0.01; SbD effect: F_(1,28)_=0.11, p=0.74, ELS by SbD interaction: F_(1,28)_= 7.08, p<0.05. Tukey’s comparison: ELS-CON different from Std-CON (*p<0.01). **(J)** *Suz12* mRNA in the NAc: Two-way ANOVA: ELS effect: F_(1,24)_= 26.87; p<0.0001; SbD effect: F_(1,24)_=0.15, p=0.70; ELS by SbD interaction: F_(1,24)_= 0.74, p=0.38. Tukey’s comparison: ELS-CON different from Std-CON (*p<0.05), ELS-SbD different from Std-SbD (**p<0.01). Data: Mean ± SEM.

### ELS increases levels of SUZ12 in the PFC

Methylation dynamics of H3K27, including its H3.3 variant, are catalyzed by PRC2^28, 47–50^, and involves the coordinated action of its core subunits: suppressor of Zeste 12 (SUZ12), enhancer of Zeste 2 (EZH2), and embryonic ectoderm development (EED)^28, 33, 51^. We therefore measured the mRNA expression levels of *Suz12*, *Ezh2*, and *Eed* in the PFC of adult male mice previously exposed to ELS. We observed higher *Suz12* mRNA expression in the PFC of ELS-exposed mice compared to their Std-raised counterparts (**Figure 3F-G**), while no differences in *Ezh2* or *Eed* mRNA were detected. Hence, we focused on *Suz12* mRNA for downstream experiments. Importantly, we replicated in the same mice ELS induction of *Suz12* in the NAc as we reported previously^18^.

We next measured *Suz12* mRNA expression in adult male mice previously subjected to ELS followed by SbD to establish whether the double-hit stress paradigm induces an additive effect in the PFC (**Figure 3H**). We found higher *Suz12* mRNA expression in mice exposed to ELS alone relative to Std-CON mice, but not in Std- or ELS-mice exposed to SbD (**Figure 3I**). By contrast, in the NAc, we observed higher *Suz12* mRNA expression in mice exposed to ELS-CON and ELS followed by SbD compared to their respective Std-raised controls (**Figure 3J**). Collectively, these findings suggest that ELS alters the epigenetic function of PRC2 by inducing SUZ12 in both the PFC and NAc, although with different temporal dynamics in the two brain regions.

### Expressing the VEFS domain of SUZ12 in the adolescent PFC or NAc controls behavioral responses to stress in a sex-dependent manner

We recently demonstrated that viral-mediated expression of the VEFS domain of SUZ12 (**Figure 4A**) selectively increases H3K27me1 accumulation in the adult NAc^19^, thus providing a novel means of manipulating H3K27me1 levels and examining its downstream consequences. Because ELS induces a transient increase in H3K27me1 in the PFC of female adolescent mice (**Figure 2F**), we infused neuron-specific adeno-associated virus (AAV) vectors to selectively express the VEFS domain (AAV-VEFS) plus GFP or GFP alone as a control (AAV-GFP) into the female PFC at PND21. Importantly, we confirmed that, as seen in the NAc, AAV-VEFS selectively increases VEFS levels in the PFC (**Figure 4B-C**). We next examined whether adolescent manipulation of AAV-VEFS in the PFC induces behavioral and cognitive alterations in adult females (**Figure 4D**). We found that AAV-VEFS-injected females displayed anxiety-like behaviors as shown by reduced time in the open arms of the elevated plus maze compared to GFP-injected females (**Figure 4E**). These mice also failed to show social preference for a conspecific mouse in the social preference test (**Figure 4F**).

**Figure 4.**
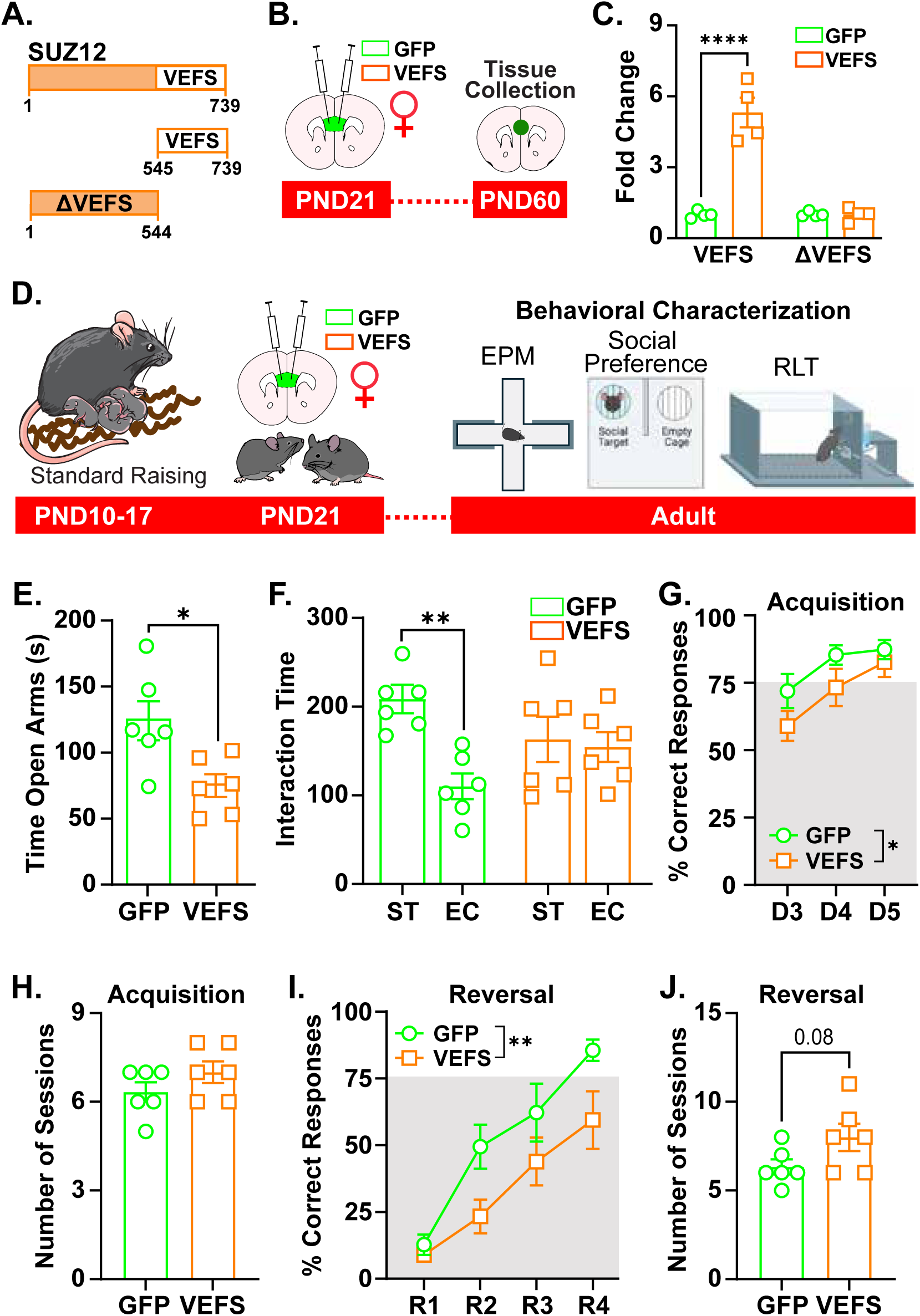
VEFS domain of SUZ12 in the adolescent PFC induces behavioral alterations in adult female mice. **(A)** Schematic of the 739-amino-acid-long SUZ12 protein. SUZ12 colored to highlight its interacting (VEFS, white) and localization (ΔVEFS, orange) domains. The VEFS domain spans 545 to 739 amino acids. **(B)** Schematic and timeline of VEFS expression in the NAc at PND21, and tissue collection in adulthood. **(C)** qPCR showing that AAV-VEFS increases levels of VEFS expression without altering levels of the N-terminal region of *Suz12* termed ΔVEFS (n=4/group). VEFS: t_(8)_=6.88, p<0.01; ΔVEFS: t_(10)_ = 0.07, p=0.94. **(D)** Timeline of VEFS-injection at PND21 and behavioral characterization in adulthood. **(E)** Time in open arms: t_(10)_=2.86, p<0.05. **(F)** Time of social interaction: Two-way ANOVA: Social Target (ST) effect: F_(1,20)_= 8.14, p=0.01; Virus effect: F_(1,20)_= 0.05, p=0.97, ST by Virus interaction: F_(1,20)_= 5.74, p<0.05. Tukey’s comparison: ST different from empty cage (EC) only in AAV-GFP-CON, **p<0.01. **(G)** Percentage of correct lever presses during acquisition: Two-way ANOVA: Session effect: F_(2,30)_=7.19, p<0.01, Virus effect: F_(1,30)_=18.32, p<0.05. Tukey’s comparison: AAV-VEFS from AAV-GFP, *p<0.05. **(H)** Number of sessions to acquisition criterion: t_(10)_=1.33, p=0.21. **(I)** Percentage of correct lever presses during reversal: Two-way ANOVA: Virus effect: F_(1,40)_=12.78, p<0.001; Session effect: F_(3,40)_= 23.51, p<0.0001. Tukey’s comparison: AAV-VEFS from AAV-GFP, *p<0.05. **(J)** Number of sessions to reversal criterion: t_(10)_=1.89, p=0.08. Data: Mean ± SEM.

We next trained AAV-VEFS- or AAV-GFP-injected female mice in the operant RLT to assess changes in cognitive flexibility. We observed a main effect of the virus, with AAV-VEFS-injected mice showing a lower percentage of correct responses during the acquisition phase of the RLT (**Figure 4G**). However, these mice did not differ in the number of trials required to reach the learning criterion (**Figure 4H**). During the reversal phase, we also observed that AAV-VEFS-injected females showed a reduced percentage of correct responses relative to AAV-GFP-injected mice (**Figure 4I**) and exhibited a trend toward an increase in the number of trials to reach the reversal criterion (**Figure 4J**). These findings are consistent with impaired cognitive flexibility induced by VEFS domain expression in the PFC of female mice. Interestingly, VEFS-induced behavioral alterations in female mice were specific to its infusion into the adolescent PFC, since the same manipulation in the female NAc at PND21 (**Figure S2A**) only led to a reduction in time in the open arms of the elevated plus maze (**Figure S2B**), but did not induce social and cognitive alterations in adulthood (**Figure S2C-E**).

We previously found that VEFS expression in D1-MSNs of the adult NAc leads to stress-induced social and cognitive alterations in male mice^19^, however, in this earlier study we did not examine the effect of VEFS expression in adolescent mice. Therefore, for completeness, we utilized Cre-dependent AAV-VEFS and AAV-GFP vectors to selectively express the VEFS domain in NAc D1-MSNs^19^. We microinfused AAV-VEFS or AAV-GFP into the NAc of D1-Cre adolescent male mice at PND21 (**Figure 5A-C**). We selected PND21 for viral injections based on our proteomic findings showing increased H3K27me1 in the NAc of ELS-exposed mice^19^. We then subjected AAV-VEFS- or AAV-GFP-injected D1-Cre mice to SbD^37^ or control conditions in adulthood (**Figure 5D**). As anticipated, AAV-GFP- and AAV-VEFS-injected control mice displayed normal social interaction towards a novel CD1 mouse (social target) (**Figure 5E**). However, AAV-VEFS-injected mice subjected to SbD showed reduced interaction time with a social target compared (**Figure 5E**), demonstrating that AAV-VEFS infusion in adolescence sensitizes susceptibility to SbD in adulthood. Notably, we also found that control and SbD mice previously infused with AAV-VEFS spent less time in the open arms of an elevated plus maze than AAV-GFP mice (**Figure 5F**), indicating an anxiogenic-like phenotype.

**Figure 5.**
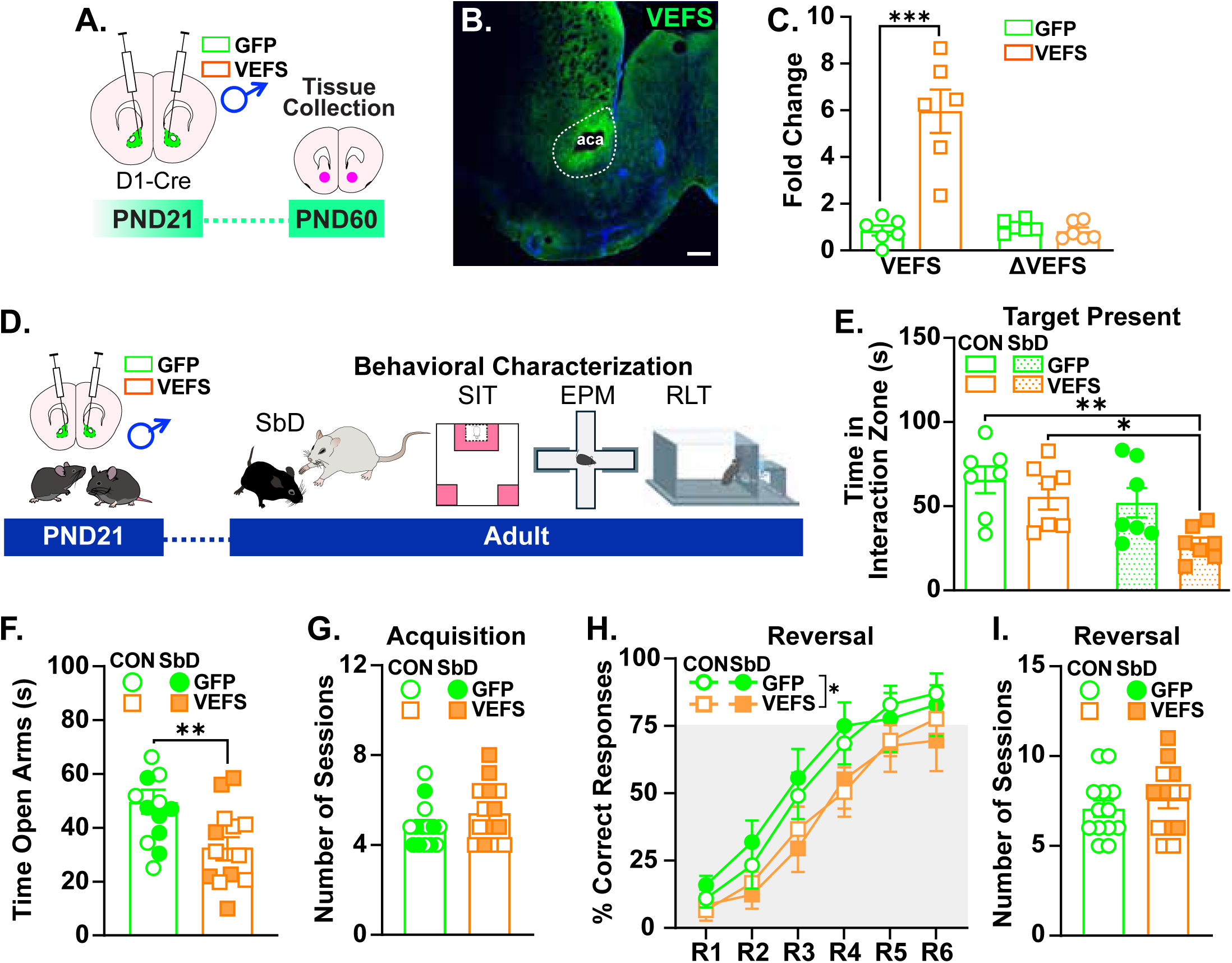
VEFS Domain of SUZ12 in the Adolescent NAc Sensitizes Stress Susceptibility in Adult Male Mice. **(A)** Schematic and timeline of VEFS expression in the NAc at PND21, and tissue collection in adulthood. **(B)** Coronal section showing GFP staining of VEFS in the adult NAc. Scale bar: 500μm. **(C)** qPCR showing that AAV-VEFS increases levels of VEFS expression without altering levels of the N-terminal region of *Suz12* termed ΔVEFS (n=6/group). VEFS: t_(10)_=5.35, p<0.001; ΔVEFS: t_(10)_ = 1.04, p=0.32. **(D)** Timeline of VEFS-injection at PND21 and exposure to SbD in adulthood. **(E)** Time of interaction with social target: Two-way ANOVA: SbD effect: F_(1,24)_= 11.90, p<0.01; virus effect: F_(1,24)_=3.002, p=0.09; SbD by virus interaction: F_(1,24)_= 4.20, p<0.05. Tukey’s comparison: VEFS-SbD different from GFP-CON (*p<0.01), VEFS-SbD different from VEFS-CON (*p<0.01). **(F)** Time in open arms: t_(23)_=2.87, p<0.01. **(G)** Number of sessions to Acquisition: t_(24)_=1.32, p=0.19. **(H)** Percentage of correct lever presses during reversal learning: Three-way ANOVA: Session effect: F_(5,144)_=44.53, p<0.0001; Virus effect: F_(1,144)_=15.12, p=0.001. Tukey’s comparison: VEFS different from GFP (*p<0.01). **(I)** Sessions to Reversal learning: t_(24)_=0.77, p=0.44. Data: Mean ± SEM.

Because male mice subjected to the double-hit stress paradigm exhibit deficits in cognitive flexibility (**Figure 1F**), we next assessed AAV-VEFS- or AAV-GFP-injected mice, exposed to SbD or control conditions, in the RLT (**Figure 5D**). As expected, there were no significant differences between groups during the acquisition phase, as all experimental mice took about the same number of sessions to achieve the learning criterion (**Figure 5G**). However, during the reversal phase, we observed that AAV-VEFS-injected mice had a lower percentage of correct responses than AAV-GFP-injected mice, an effect that was independent of adult exposure to SbD (**Figure 5H**) but did not exhibit an increase in the number of trials to reach the reversal criterion (**Figure 5I**). Overall, these findings indicate that VEFS expression in the adolescent NAc of males, but not in females, leads to behavioral and cognitive alterations that can be exacerbated by adult exposure to stress.

## DISCUSSION

Adverse experiences during early life can render the brain more reactive to future environmental insults^2, 3^, and sensitize stress responses long after the initial stressors have ended^17–19, 23, 25^. The effects of stress are known to influence transcriptional and epigenetic changes in a sex-specific manner, leading to divergent behavioral outcomes across the lifespan^52–56^. Therefore, gaining a better understanding of the sex differences that influence the lasting changes induced by ELS at the molecular and circuit levels is crucial. This understanding could facilitate the development of new therapeutic and preventive strategies for vulnerable individuals to stress^1,15, 17, 21^.

Our prior study of adult NAc was the first to establish a novel function of H3K27me1 in the brain, and specifically implicate it as a key regulator of lifelong stress susceptibility^19^. Using unbiased histone profiling and genome-wide sequencing, we showed that male, but not female, mice subjected to ELS displayed a lasting increase in H3K27me1 abundance in the NAc^19^. This increase was further amplified by a subsequent exposure to SbD in adulthood^19^. In the present study, we found that ELS also induces H3K27me1 accumulation in the PFC, but with distinct sex-specific and age-dependent features as seen for NAc. For example, we found that male mice subjected to ELS exhibited H3K27me1 accumulation in the PFC only in adulthood, whereas ELS-exposed females exhibited higher H3K27me1 abundance in this brain region only during adolescence. We also found that H3K27me1 accumulation in response to ELS was associated with mRNA induction of the *Suz12* subunit of PRC2 in both the PFC and NAc, supporting the strong connectivity between these two regions in mediating the lasting effects of stress^57–59^. Indeed, we show here that the region-specific expression of the VEFS domain of SUZ12, which induces H3K27me1 levels, during adolescence led to sex-specific social and cognitive alterations in adulthood, mimicking, at least in part, some of the behavioral alterations induced by ELS. Together, these findings indicate that H3K27me1 is an important chromatin scar across the corticolimbic pathway that mediates the sex-specific behavioral effects of ELS.

The role of H3K27me1 in the brain has been poorly studied despite its higher abundance compared to other histone modifications, such as H3K27me3^60^, and little is known about the mechanism by which H3K27me1 influences male and female transcriptional states, either across development or in response to environmental insults. Our histone profiling studies revealed that H3K27me1 accumulation is differentially regulated in the PFC and NAc under baseline conditions. For instance, the abundance of H3K27me1 in the PFC does not change between adolescence and adulthood in Std-raised male and female mice (**Figure 2D-F**). By contrast, in the NAc, only males show high H3K27me1 abundance during adolescence but very low levels in adulthood^19^, a pattern that is disrupted by ELS exposure^19^. We also showed previously that H3K27me1 binding to DNA in the NAc occurs across genes that control neuronal excitability, such as ionotropic receptors and voltage-gated channels, and synaptic plasticity, including axonal guidance cues or cell adhesion molecules^19^. These findings suggest that ELS may differentially disrupt H3K27me1-dependent epigenetic programs in the PFC and NAc, leading to distinct transcriptional responses that vary in an age- and sex-dependent manner and ultimately impair the functional connectivity between these two regions^57, 58, 61^. Future experiments will be critical to determine whether genomic enrichment of H3K27me1 in the PFC occurs at a similar set of genes to those bound in the NAc and identify the precise role of the PRC2 machinery in modulating ELS-induced H3K27me1 enrichment at a genome-wide level. This information will be critical for the use of more advanced molecular techniques, including viral-mediated CRISPR-Cas9 systems^62, 63^, to functionally manipulate H3K27me1 enrichment at specific genomic loci.

Our previous work has shown that ELS shapes the transcriptional profiles of the brain’s reward circuitry, including the PFC and NAc^17, 18, 25^. This transcriptional regulation is mediated, in part, by enrichment of H3K79me2 in D2-MSNs of the NAc, and the coordinated action of its writer, DOT1L, and eraser, KDM2B, enzymes^18^. Here, we found that ELS led to H3K27me1 changes that predominate in pyramidal neurons of the PFC (**Figure 3B-C**) as well as in D1-MSNs (and perhaps D2-MSNs to a lesser degree) of the NAc (**Figure 3D-E**). These results indicate that ELS may facilitate the epigenetic crosstalk between histone modifications^64^, thus, promoting the enrichment or depletion of other histone marks. Such a differential pattern of histone modifications may be influenced by the degree of neuronal activation following ELS exposure^22, 65, 66^, as well as the strength of connectivity between brain regions^22, 65, 66^. Indeed, recent evidence revealed that neurons of the NAc that are activated by ELS exhibited dynamic changes of chromatin accessibility^22^, an effect that was linked to H3K4me1^22^, a key epigenetic modification for active enhancers^67^. In future studies, we will explore the contribution of ELS-induced H3K27me1 in altering the epigenetic landscape across cell types of the PFC and NAc.

Our previous work on adult exposure to chronic social defeat stress (CSDS) revealed a strong correlation between social interaction and performance in the RLT, with susceptible mice after CSDS displaying reduced cognitive flexibility^19^. Here we showed that male and female mice exposed to ELS followed by SbD exhibited high social avoidance compared to Std-raised mice or mice subjected ELS alone (**Figure 1B-C**), which is consistent with our previous ELS studies^17–19^. We also found that ELS led to sex-specific alterations in cognitive flexibility in the RLT: while ELS, in combination with SbD, impaired reversal learning in males (**Figure 1E**), ELS affected reversal learning in females regardless of adult SbD exposure (**Figure 1F**). However, our results also showed that male and female mice differed in how ELS affected cognitive flexibility, for example, males were impaired during late stages of the RLT, whereas females exhibited impaired performance during early stages. Evidence shows that increased perseveration or decreased consolidation of novel strategies are core features of inflexible behavior^68–71^, yet these cognitive alterations involve distinct patterns of corticolimbic activity^68–71^. In this context, we hypothesize that ELS alters the functional connectivity of the corticolimbic pathway that results in sex-specific alterations in reversal learning. Our finding that viral-mediated VEFS expression in the adolescent PFC (in females) or NAc (in males) led to sex-specific social and cognitive alterations in adult mice warrants future studies aimed at understanding the role of H3K27me1 in mediating the molecular and functional adaptations across the corticolimbic pathway following exposure to ELS.

## ACKNOWLEDGEMENTS

This work was supposed by grants from NIMH (R01MH129306) and the Hope for Depression Research Foundation to E.J.N.; the Robin Chemers Neustein Award, the FBI Innovation Award, and the Mass General for Children Microgrant to A.T.B.; the Lurie Center Summer Research Program to N.B, and the Harvard College Research Program to N.A. The Sidoli lab gratefully acknowledges for funding the Hevolution Foundation (AFAR), the ERC-CFAR Center for AIDS research, the Einstein-Mount Sinai Diabetes center, and the NIH Office of the Director (S10OD030286). This study used the equipment of the Boston Area Nutrition Obesity Research Center supported by NIH (DK040561), the Microscopy and Advanced Bioimaging CoRE at Mount Sinai and the Lurie Center for Autism Research Core.

Illustrations were designed by Jill Gregory (Mount Sinai), Carlos Torres-Berrío or by created using BioRender.com templates.

## AUTHOR CONTRIBUTIONS

A.T.B. and E.J.N. designed the studies. A.T.B., J.N., N.B., and N.A. performed stereotaxic surgeries, tissue collection behavioral, molecular, and neuroanatomical experiments. A.B. helped with confocal imaging. S.S. conducted proteomics experiments. S.J.R. helped with female defeat experiments. A.T.B. and E.J.N. supervised the project and wrote the manuscript. All authors discussed the results and commented on and edited the manuscript.

## MATERIALS AND METHODS

### Animals

Experimental procedures were performed in accordance with the guidelines of the Institutional Animal Care and Use Committee (IACUC) at the Icahn School of Medicine at Mount Sinai and the Massachusetts General Hospital. All male and female mice used in these studies were maintained on a 12-hour light-dark cycle (lights on at 7:00) with ad libitum access to food and water throughout the experiments. All data derived from animal studies were analyzed by an experimenter blind to the conditions.

Male and female C57BL/6J wild-type mice (Postnatal day, PND, 75±15, Jackson Laboratory) served as experimental subjects in the social defeat stress paradigms. Male and female mice were group-housed before exposure to the stress procedures and single-housed at the completion of the last defeat session and before the social interaction test (SIT). Animals were randomized by cage prior to exposure to stress or before stereotaxic surgeries (e.g., each cage was assigned to each experimental condition or treatment: control or manipulation). The order of the animals was further randomized prior to behavioral tests.

Male CD-1 retired breeder mice (≥3 months old) previously screened for aggressive behavior were used as social aggressors for male stress exposure. CD-1 mice were obtained from Charles River Laboratories and were single-housed throughout the study. Male ERα-Cre (Line: 017911, B6N.129S6(Cg)-Esr1tm1.1(cre)And/J0 were obtained from Jackson laboratory, and were crossed with CD-1 females to obtain F1 males, which were used as aggressors for female CSDS^72^. For female SbD, ERα-Cre F1 mice infused with either mCherry or hM3D(Gq)-mCherry were injected intraperitoneally with 1DmgDkg^−1^ of clozapine N-oxide (CNO, Tocris Bioscience) 20 minutes before each defeat session. Male *Drd1a*-Cre hemizygote bacterial artificial chromosome transgenic mice on a C57BL/6J background (Line FK150, http://www.gensat.org/cre.jsp) were bred in our colony room. *Drd1a*-Cre mice were group-housed until the beginning of social defeat experiments and single-housed at the completion of the last defeat session and before all behavioral testing and downstream experiments.

### Stress Paradigms and Behavioral Tests

<u>Early life stress (ELS):</u> Two adult C57/BL6J wild-type female mice (PND 75±15) were mated with one adult male mouse from the same strain in our animal facilities. The male was removed after 1 week, and the females were separated into individual clean cages 2–3 days before the expected date of delivery (PND0). Litters were weighed and counted at birth and on PND7, and cages were cleaned on PND10, but were otherwise undisturbed. Only litters with 5 to 9 pups were included in this study, and the litter size was balanced across ELS and standard-raised conditions (Std).

The ELS paradigm consisted of a combination of both maternal separation, in which the entire litter was removed from the dam to a clean cage for 4 hours/day, and nesting material (crinkled Enviro-Dri paper) was reduced to one-fourth the regular amount during PND10 to PND17. We selected this developmental period based on our previous research which demonstrated enhanced susceptibility in adult mice exposed to this ELS paradigm without any overt behavioral abnormalities in the absence of this second hit of adult stress^17, 18, 25^. Upon completion of the ELS protocol, nesting material was restored at PND17 and pups stayed with their dams until weaning at PND21. Std pups were monitored daily from PND10 to PND17, but cages contained regular amount of nesting material and pups were not separated from their dams.

<u>Subthreshold social defeat (SbD):</u> SbD was performed as in^37, 43^ and consisted of three sessions of social defeat occurring on a single day. Briefly, each adult male C57BL/6J experimental mouse was exposed to 5 minutes of physical aggression by a male CD-1 mouse. At the completion of the session, C57BL/6J experimental and CD-1 mice were housed in a 2-compartment hamster cage and separated by a transparent divider with holes to provide sensory, but not physical, contact for 15 minutes, followed by two additional defeat exposures. After the SbD sessions, C57BL/6J experimental mice were single-housed for 24 hours prior to the SIT.

Control C57BL/6J mice were housed in 2-compartment cages with a cage-mate during the same period. The SbD protocol was conducted during the light cycle, between 11:00 and 14:00.

<u>Female SbD</u>: SbD was performed as in^19, 72^. In brief, females were exposed to 5 minutes of physical aggression by a male ERα-Cre mouse and returned to their home cage for 15 minutes, followed by two additional defeat exposures. During the SbD, females were group-housed but single housed after the last defeat session.

<u>Social interaction test (SIT):</u> Twenty-four hours after the last session of SbD, C57BL/6J experimental mice were assessed in the SIT. This test consisted of 2 sessions in which defeated and control mice explored a squared-arena (44 x 44 cm) in the absence or presence of a novel aggressor CD-1 mouse (social target) for a period of 2.5 minutes each session. In the first session, an empty wire mesh enclosure 10 cm (length) × 6.5 cm (width) × 42 cm (height) was located against one of the walls of the arena to assess baseline exploration. In the second session, an unfamiliar CD-1 aggressor was placed inside the wire mesh enclosure. The area that surrounded the enclosure was designated as the social interaction zone (14 cm x 9 cm), whereas the corner areas of the walls opposite to the enclosure were designated as corners (9 x 9 cm) and represented the farthest point from the social interaction zone. The time (in seconds) of interaction with the social target was measured during both sessions as in^37^. This simple measure has been shown to correlate strongly with numerous other behavioral outcome measures^19^.

<u>Elevated plus maze:</u> Anxiety-like behavior was assessed in a plus maze elevated 50 cm from the floor. The maze consisted of two facing open arms and two facing enclosed arms that extend from a central platform. Mice were placed in the center platform of the maze facing one of the open arms and left to explore the arms during five minutes. The time spent in the open versus closed arms of the maze were recorded with an overhead video camera and analyzed using the EthoVision XT 11 (Noldus Leesburg).

<u>Sociability test:</u> Sociability to a novel conspecific mouse was assessed in an open field (44 x 44 cm) which containing two enclosures placed in opposite corners of the arena. The area surrounding the enclosures (19 cm x 23 cm) was designated as interaction quadrants. During habituation, each experimental mouse was allowed to freely explore the open field and empty enclosures for 8 minutes. For the sociability test, a juvenile male conspecific (∼4 weeks old) -a social target- was placed in one of the enclosures, while the other enclosure remained empty. The experimental mouse was allowed to explore the open field containing a social target on one side and an empty enclosure on the other side for 8 minutes. The time spent in each quadrant with the social target and the empty enclosure were recorded.

<u>Reversal learning task:</u> Reversal learning was conducted in mouse operant chambers (Interior dimensions: Interior: 55.69 cm x 38.1 cm x 40.64 cm; exterior dimensions: 63.5 cm x 43.18 cm x 44.45 cm, and walls: 1.9 cm from Med Associates (St. Albans). Operant chambers were enclosed in light and sound attenuating cubicles equipped with white house lights as well as fans to provide ventilation and to mask external noise. Each operant chamber contained two retractable levers, located on the right and left sides of a central reward magazine calibrated to deliver ∼50 µl of liquid. Adult male mice were water deprived and given 4 hours of water access during the 3 days prior to the beginning of behavioral training. They received a single operant session every day and were given 2 hours of water access following each daily session throughout the course of the experiment. During the single 30 minutes pre-training session, mice explored the operant chamber and learned to introduce their noses into the central reward magazine to get 0.2% saccharin rewards, which were delivered every 60 seconds. Levers were retracted throughout this pre-training.

The reversal learning task consisted of acquisition and reversal phases. The acquisition phase was further divided into “ad libitum” and “restricted” sessions. The “ad libitum” sessions were given during the first three days of the acquisition phase and consisted of 30 minutes of free access to the active (correct) and inactive (incorrect) levers. Mice were allowed to freely press each lever, but only the active lever granted access to 10 seconds of 0.2% saccharin-water reward at a fixed-ratio 1 (FR1) schedule. During the “restricted” sessions, the active and inactive levers were available but mice had only 30 seconds to make a choice. Levers were immediately retracted once the choice was made and the reward was delivered if the correct lever was pressed. If mice did not press any of the levers during the 30 second trial or pressed the incorrect lever, they entered a 30 second timeout and an omission or an incorrect response was recorded. The number of correct and incorrect lever presses along with the number of earned rewards and response omissions were measured for each session, with a learning criterion of 75% correct presses for three consecutive days. During the reversal phase, only “restricted” sessions were given, in which the correct lever was switched, thus, all experimental mice would need to learn to press the previously inactive lever to obtain the 0.2% saccharin reward. The number of correct and incorrect lever presses along with the response omissions and the number of earned rewards were recorded, and a learning criterion of 75% correct presses for three consecutive days was used.

### Tissue Dissection

Experimental mice were euthanized by rapid decapitation. Brains were removed, and cooled with ice-cold PBS prior to slicing on a pre-defined brain matrix. Unilateral 12-gauge punches of the PFC and bilateral 14-gauge punches of the NAc were taken from 1 mm coronal sections starting on approximately plate 14 or 15 of the Paxinos & Franklin mouse atlas, frozen immediately on dry ice, and stored at −80°C until further use. For ELS experiments, tissue was collected at PND21 and adulthood, after PND60. Bilateral 15-gauge punches were used for NAc in PND21 mice.

### Mass Spectrometry

<u>Nuclei isolation and histone purification:</u> Frozen tissue from the NAc was washed in a 2 ml pre-chilled dounce homogenizer containing 500 µl of nuclear isolation buffer (15 mM Tris, 60 mM KCl, 15 mM NaCl, 5 mM MgCl_2_, 1 mM CaCl_2_, 250 mM sucrose) with protease inhibitors and stabilizing agents (1 µl of 1 M DTT, 2.5 µl of 200 mM 4-benzenesulfonyl fluoride hydrochloride, AEBSF, 2 µl of 2.5 µM microcystin and 2 µl of 5 M sodium butyrate, for each 1 ml), and dissociated on ice in 500 µl of lysis buffer (nuclear isolation buffer + 0.2% IGEPAL) as previously described^39^. For histone purification, nuclei were incubated in 0.2 M H_2_SO_4_ for 2 hours with constant rotation at 4°C, followed by centrifugation at 3,400 rcf for 5 minutes. Collected supernatant was added with chilled 33% trichloroacetic acid (Sigma-Aldrich), and left on ice at 4°C for 1 hour. After centrifugation at 3,400 rcf for 5 minutes, the supernatant was discarded and the histone pellet was rinsed first with 150 µl of ice-cold acetone + 0.1% HCl, and then with 150 µl of 100% ice-cold acetone. Histones were allowed to air-dry for 5 minutes, dissolved in 20 µl ddH_2_O and stored at −80°C for further applications.

<u>Histone derivatization and digestion:</u> Purified histones were treated with 20 μl of 50 mM NH_4_HCO_3_ (pH 8.0), and split into two aliquots for digestion into ArgC-like peptides (bottom-up) and intact histone N-terminal tails (middle-down). For the bottom-up preparation, samples were treated with 5 µl of acetonitrile followed by 5 µl of propionic anhydride and 14 µl of ammonium hydroxide and incubated for 20Dminutes. Samples were then dried, resuspended in 20 µl of 50 mM NH_4_HCO_3_ and digested with Trypsin (Promega) at an enzyme:sample ratio 1:20 for 2 hours. The derivatization reaction was then performed again twice to derivatize peptide N-termini. Samples were then desalted by using in-house packed C18 Stage-tips and dried using a SpeedVac centrifuge prior to Liquid Chromatography–Mass Spectrometry (LC-MS/MS) analysis.

<u>LC-MS/MS:</u> Samples were resuspended in 10 μl of water + 0.1% formic acid. A volume of 2 μl of histone peptide solution was injected onto a 75 µm ID x 25 cm Reprosil-Pur C18-AQ (Dr. Maisch Beim Brückle) nano-column packed in-house. The LC-MS setup consisted in a Dionex RSLC Ultimate 3000 (Thermo Scientific), coupled online with an Orbitrap Fusion Lumos (Thermo Scientific). The HPLC gradient was as follows: 2% to 28% solvent B (AD=D0.1% formic acid; BD=D95% MeCN, 0.1% formic acid) over 45Dminutes, from 28% to 80% solvent B in 5Dminutes, 80% B for 10Dminutes at a flow rate of 300Dnl/minute. The mass spectrometer was set to acquire spectra in a data-independent acquisition (DIA) mode. Briefly, the full MS scan was set to 300-1100 m/z in the orbitrap with a resolution of 120,000 (at 200 m/z) and an AGC target of 5×10e5. MS/MS was performed in the orbitrap with sequential isolation windows of 50 m/z with an AGC target of 2×10e5 and an HCD collision energy of 30 as described in^39^. Extraction of the signal of the (un)modified peptides was performed by using EpiProfile 2.0, a computational platform that allows the accurate quantification of histone marks^73^. From the extracted ion chromatogram, the area under the curve was obtained and used to estimate the abundance of each peptide. To achieve the relative abundance of post-translational modifications (PTMs), the sum of all different modified forms of a histone peptide was considered as 100% and the area of the particular peptide was divided by the total area for that histone peptide in all of its modified forms. The relative ratio of two isobaric forms was estimated by averaging the ratio for each fragment ion with different mass between the two species. The resulting peptide lists generated by EpiProfile 2.0 were exported to Microsoft Excel and further processed for a detailed analysis.

### Neuroanatomical Experiments with Mouse Brain Tissue

<u>Immunofluorescence:</u> Mice were anesthetized with a lethal dose of pentobarbital sodium (Vortech), and perfused intracardially with 0.9% saline, followed by 4% paraformaldehyde (PFA) in PBS (pH 7.4). Brains were removed and maintained in 4% PFA overnight. Coronal sections of the NAc were obtained at 30 µm using a vibratome (Leica VT1000S). For double-labeled immunofluorescence, PFC or NAc sections were blocked with 2% BSA + 0.2% Tween 20, and incubated overnight at 4°C with anti-H3K27me1 (1:500), anti-SMI-32 antibodies (1:500), or Chicken anti-GFP antibody (1:1000). Immunostaining was visualized with Alexa 488-conjugated (Jackson Immunoresearch), and Cy3-conjugated (Jackson Immunoresearch) secondary antibodies raised in donkey. RNAscope images were taken within 2 weeks of staining on a Zeiss LSM780 confocal microscope and immunofluorescence images were taken on an Andor BC43 confocal microscope. All images were analyzed using a macro from ImageJ^74^ to find intensity of SMI-32 or H3K27me1 staining for each DAPI-positive nucleus. Images were captured at 512 × 512 pixels, and 140 × 140-pixel. Nuclei above the 75th percentile of SMI-32 expression were deemed SMI-32. Oversaturated images of H3K27me1 staining were excluded from the analysis. The percentage of change in average H3K27me1 intensity was calculated for a total of four animals per group with at least two bilateral sections analyzed per animal.

<u>RNAscope and immunofluorescence:</u> Combined RNAscope and immunofluorescence staining was done as in^19^. Briefly, mice were anesthetized with a lethal dose of pentobarbital sodium, and perfused intracardially with 50 ml of 0.9% saline, followed by 75 ml of ice-cold fixative solution (4% PFA in PBS; pH 7.4). Brains were immersed in 10%, 20% and 30% sucrose in 0.2 M PB at 4°C until sunk and flash frozen with cold 2-methylbutane (Fisher Scientific). Coronal sections of the NAc, spanning plates 15–24 of the Paxinos & Franklin mouse atlas, were obtained at 30 μm using a cryostat (Leica CM3050 S), mounted onto SuperFrost Plus (Fisher Scientific), and maintained in the −80°C freezer until processing. For immunofluorescence followed by RNAscope, tissue was processed using the Multiplex Fluorescent Reagent Kit v2 protocol (Advanced Cell Diagnostics), with minor modifications. In brief, brain sections were first acclimatized at −20°C for ∼2 hours, and then at RT° for ∼1 hour. Sections were incubated in 4% PFA for 30 minutes, rinsed three times with PBS and air-dried for at least 15 minutes. Slices were then baked in the HybEZ Oven at 60°C for 30 minutes and dehydrated with xylene followed by ethanol 100%. Dried sections were treated with RNAscope hydrogen peroxide, and Co-Detection Target Retrieval Solution at 100°C for 15 minutes (Advanced Cell Diagnostics). After several washes with distilled water and PBS, sections were incubated overnight at 4°C in anti-H3K27me1 (1:300) antibody diluted in Co-Detection Antibody Diluent. For RNAscope, tissue was washed and incubated in RNAscope Protease plus at 40°C for 30 minutes in the HybEZ Oven, followed by hybridization in Mm-*Drd1a*, and in Mm-*Drd2* probes to label *Drd1* and *Drd2* MSNs, respectively. Slides were counterstained with DAPI for 5 minutes and cover-slipped with ProLong Diamond Antifade Mountant (Thermo Scientific). Images were taken within 2 weeks of staining on a Zeiss LSM780 confocal microscope and analyzed using a macro from ImageJ^74^ to find intensity of *Drd1*, *Drd2*, or H3K27me1 staining for each DAPI-positive nucleus. Images were captured at 512 × 512 pixels, and 140 × 140-pixel. Nuclei above the 75th percentile of *Drd1* expression were deemed *Drd1*+ and likewise for *Drd2*+ nuclei. Oversaturated images were excluded from the analysis. The percentage of change in average H3K27me1 intensity was calculated for a total of four animals per group with at least three bilateral sections analyzed per animal.

### RNA Extraction and Quantitative Real-Time PCR

Total RNA was isolated from mouse frozen tissue with the miRNeasy Micro Kit protocol with minor modifications (Qiagen). All RNA samples were determined to have 260/280 and 260/230 values ≥1.8, using the NanoDrop One C system (Thermo Scientific). RNA integrity was assayed using an Agilent 2100 Bioanalyzer (Agilent). Reverse transcription was performed using High-Capacity cDNA Reverse Transcription kit (Applied Biosystems). Real-time PCR was carried out with SYBR green (Applied Biosystems) using an Applied Biosystems QuantStudio 5 Pro Real-Time PCR System. Data from target genes was analyzed by comparing C(t) values using the ΔΔC(t) method and *Gapdh* was used as reference gene. Real time PCR was run in technical triplicates.

### Plasmids and Viral Constructs

Plasmids of full-length *Suz12* and of the VEFS-BOX domain (VEFS), corresponding to the C-terminal portion of SUZ12, were kindly donated by Kristian Helin (Institute of Cancer Research, UK). The full-length *Suz12* plasmid spans residues 1 to 739 and the VEFS plasmid spans residues 545 to 739 in the encoded proteins, as described in^29, 31^. Viral constructs were obtained from the Virus Vector Core (VVC) of the University of Maryland. Adeno-associated virus (AAV9) expressing a Cre-dependent VEFS-eGFP (6.89 x 10^11^ vg/ml) or SUZ12-eGFP (2.9 x 10^12^ vg/ml) fusion protein under the control of the human synapsin 1 (hSYN) promoter were used to overexpress SUZ12 or VEFS in D1-MSNs selectively. A Cre-dependent construct fused to hSYN-eGFP (9.8 x 10^11^ vg/ml) was used as control virus. Validation of the viral constructs was achieved via real time PCR using primers that recognize either the C-terminus domain of SUZ12 (VEFS) or the N-terminus domain (ΔVEFS). For female SbD, the excitatory DREADDs pAAV-hSyn-DIO-hM3D(Gq)-mCherry (2.0×10^12^ vg/ml, Plasmid #44361, Addgene) was used.

### Stereotaxic Surgery

All surgeries were performed under aseptic conditions. Adult male D1-Cre mice were deeply anesthetized with an intraperitoneal injection of rodent cocktail (ketamine: 100 mg/kg and xylazine: 10 mg/kg, diluted in 0.9% saline) and placed in a stereotaxic apparatus (Kopf Instruments). Bilateral microinfusions were made using stainless-steel infusion cannulae (33 gauge) into the mouse NAc at the following coordinates: +1.6 mm (A/P), ±1.5 mm (M/L), −4.4 mm (D/V), and 10° angle relative to Bregma. For GFP, SUZ12 and VEFS overexpression experiments, a total volume of 0.5 µl of virus was delivered on each hemisphere over a 5 minutes period. The infusion cannulae were left inside the brain area during a 5 minutes pause to prevent virus reuptake. Mice recovered for 21 days before social defeat paradigms when transgene expression is maximal. For experiments involving DREADD-mediated inhibition, a total volume of 0.5 µl of either mCherry or hM4D(Gi)-mCherry viruses was injected into the NAc of each hemisphere, as described above. For aggressors used in female CSDS, ERα-Cre F1 mice were infused bilaterally with 0.5 µl of either mCherry or hM3D(Gq)-mCherry viruses into the ventromedial hypothalamus at the following coordinates: −1.5 mm (A/P), ±D0.7 mm (M/L), −5.7Dmm (D/V) from bregma. At the completion of the behavioral testing, experimental mice were euthanized for neuroanatomy and molecular experiments.

### Quantification and Statistical Analysis

Sample sizes (n) and statistical tests used are indicated in the figure legends. All values were represented mean ± S.E.M. Statistical analysis was performed using Graphpad Prism 6.0. A significance threshold of α<0.05 was used in all the experiments. Statistical differences between two groups were analyzed with Student’s t-tests with two-tailed analysis. Correlations were calculated using the Pearson correlation coefficient. Otherwise, one-way, two-way or three-way ANOVAs were performed, followed by Tukey’s or Sidak’s multiple comparison tests. Outliers were screened using the ROUT method. No statistical methods were used to determine the sample sizes, but the number of experimental subjects is similar to sample sizes routinely used in our laboratory and in the field for similar experiments. Data with normal distribution, and similar variance were analyzed with parametric statistics. Otherwise, nonparametric statistics were applied.

## SUPPLEMENTARY INFORMATION

**Figure S1. Relative abundance of histone modifications in the PFC after ELS.** Related to Figure 2. **(A)** Heatmap displays of the relative abundance obtained for the modifications shown on H3 lysine (K) residues in standard-raised (Std), and early life stress (ELS) exposed male and female mice (n=5 samples from 3-4 different litters, 2 mice/group pooled tissue). Mono/di/trimethylation – me1/2/3; acetylation – ac. PND: postnatal day. Three-way ANOVA of each mark was calculated. Tukey’s test: ‡different from Std, p<0.0001; **different from Std, p<0.01.

**Figure S2. VEFS domain of SUZ12 in the adolescent NAc and behavioral assessment in adulthood**. **(A)** Timeline of VEFS-injection at PND21 and behavioral characterization in adulthood. **(B)** Time in open arms: t_(14)_=2.72, p<0.05. **(C)** Time of social interaction: Two-way ANOVA: Social Target (ST) effect: F_(1,20)_= 34.18, p<0.0001; Virus effect: F_(1,20)_= 0.17, p=0.68, ST by Virus interaction: F_(1,20)_= 0.04, p=0.83. Tukey’s comparison: ST different from empty cage (EC), **p<0.01. **(D)** Percentage of correct lever presses during acquisition: Two-way ANOVA: Session effect: F_(2,42)_=8.04, p<0.01; Virus effect: F_(1,42)_=1.15, p=0.28; Session by Virus interaction: F_(2,42)_=1.76, p=0.18. Tukey’s comparison: D4 different from D5, *p<0.05. **(E)** Percentage of correct lever presses during reversal: Two-way ANOVA: Virus effect: F_(1,70)_=0.05, p=0.81; Session effect: F_(4,70)_=15.20, p<0.0001. Session by Virus interaction: F_(4,70)_=0.011, p=0.99. Tukey’s comparison: R1 different from R3, R4 and R5, *p<0.05.

**Table S1.**

Raw data of single posttranslational modifications in male and female mice exposed to ELS. Related to Figure 2.

