## Supplementary figures and images for "Sex-Specific Induction of H3K27me1 in the Prefrontal Cortex Mediates the Enduring Effects of Early Life Stress"

### Supplemental File

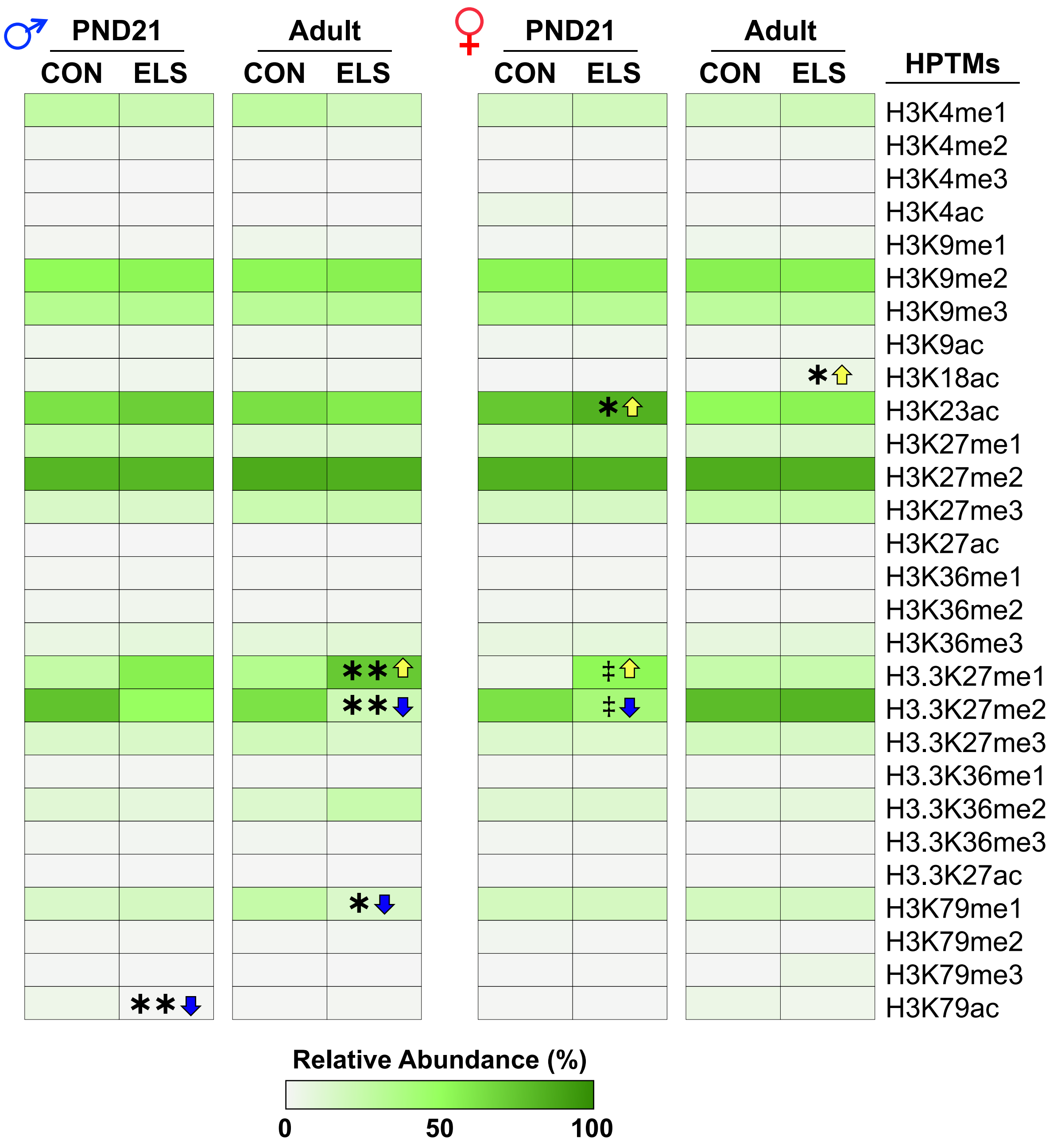

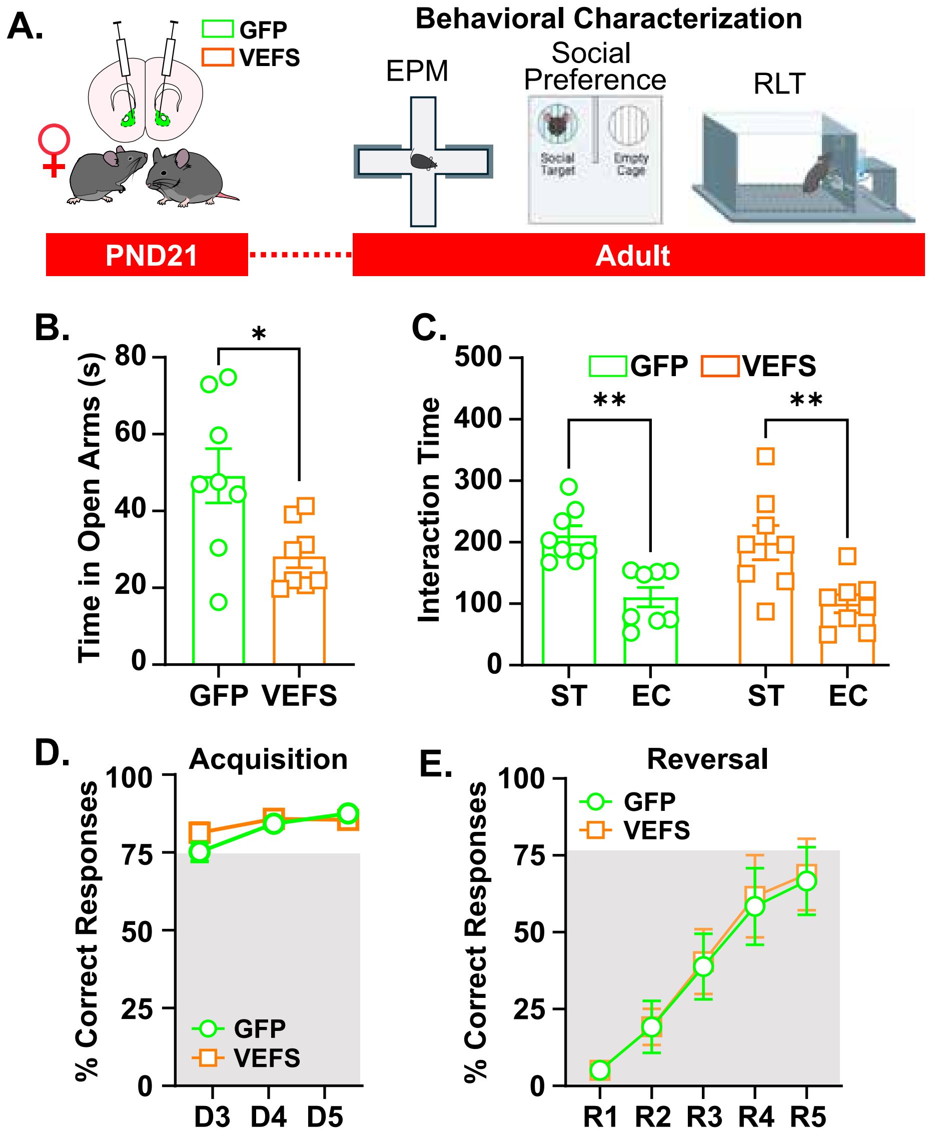
